# Density-guided AlphaFold3 uncovers unmodelled conformations in *β*_2_-microglobulin

**DOI:** 10.64898/2026.02.27.708490

**Authors:** Advaith Maddipatla, Sanketh Vedula, Alex M. Bronstein, Ailie Marx

## Abstract

Although X-ray crystallography captures the ensemble of conformations present within the crystal lattice, models typically depict only the most dominant conformation, obscuring the existence of alternative states. Applying the electron density-guided AlphaFold3 approach to *β*_2_-Microglobulin highlights how ensembles of alternate backbone conformations can be systematically modeled directly from crystallographic maps. This study also highlights how the detection of conformational ensembles is affected by the local quality of electron density and subtle variations in crystallization conditions and lattice packing. These results demonstrate that density-guided AlphaFold3 can uncover conformational heterogeneity missed by conventional refinement, offering a robust, systematic framework to capture the full structural landscape of proteins in crystals and enhancing the interpretive power of macromolecular crystallography.

**Synopsis:** Electron-density-guided AlphaFold3 reveals previously unmodeled conformational heterogeneity in *β*_2_-Microglobulin and shows how crystal packing influences ensemble detection in X-ray crystallography.

## 1 Introduction

X-ray crystallography has been the cornerstone of macromolecular structure determination for more than half a century and remains one of the most widely used techniques for resolving protein conformations at atomic resolution (Smyth & Martin, 2000). Although often interpreted as yielding a single, static representation of a biomolecule, crystallographic diffraction fundamentally reports on the ensemble of conformations present within the crystal lattice. Each Bragg reflection encodes a spatial and temporal average over billions of unit cells, and the resulting electron density map reflects the superposition of all conformational states that satisfy both the intrinsic flexibility of the macromolecule and the constraints of lattice packing. In principle, this ensemble nature should allow the characterization of conformational heterogeneity, local disorder, and transient structural states that coexist within the crystalline environment (Smith *et al*., 1986; Furnham *et al*., 2006). Despite this intrinsic ensemble averaging, macromolecular crystallography has traditionally been analyzed through the lens of a single, dominant conformation and conformational diversity is frequently under-represented, even when supported in the electron density.

Dual or multiple conformations provide a middle ground between a fully single-state interpretation and the complex, often intractable, modeling of continuous heterogeneity (van den Bedem & Fraser, 2015; Wankowicz *et al*., 2024; Riley *et al*., 2021). Alternative side-chain rotamers (Shapovalov & Dunbrack, 2011), alternative backbone traces (Rosenberg *et al*., 2024), and distinct loop or domain arrangements can be modeled explicitly when supported by residual or difference electron density. However, dual conformations remain markedly under-modeled relative to their likely prevalence(Rosenberg *et al*., 2024). Several factors contribute to this: (i) traditional refinement packages historically required manual intervention to introduce alternative conformers; (ii) avoidance of overfitting; (iii) the inherently weaker electron density corresponding to minor populations; and (iv) reporting conventions have not always encouraged detailed representation of structural heterogeneity.

We have recently developed *experiment-guided AlphaFold* (Maddipatla *et al*., 2025a; Maddipatla *et al*., 2025b; Maddipatla *et al*., 2024), a method that provides an efficient and systematic means of explicit modeling of conformational ensembles by treating AlphaFold3 (Abramson *et al*., 2024) as a structural prior. This methodology also allows reanalysis of electron density maps for structures already deposited to the protein data bank (PDB) (Joosten *et al*., 2014; Burley *et al*., 2017). Here we demonstrate the usefulness of this method on the *β*_2_-microglobulin (*β*_2_M) protein for which large amounts of structural data are available in the PDB and demonstrate that our method uncovers previously unmodelled alternate backbone conformations whose inclusion in the model provide a better fit to the electron density.

*β*_2_M is a small protein that adopts a classical immunoglobulin (Ig)-like fold, consisting of seven antiparallel *β*-strands arranged into two *β*-sheets that form a compact *β*-sandwich. In major histocompatibility complex (MHC) class I molecules, *β*_2_M binds with the heavy chain, stabilizing the overall fold and indirectly supporting peptide binding in the antigen-presenting groove (Esposito *et al*., 2005) (Figure 1, left). A key structural feature is a binding interface loop harboring a highly conserved tryptophan residue (`W60` in human *β*_2_M). Two out of the tens of high-resolution crystal structures for the isolated *β*_2_M have revealed alternate backbone conformations in this loop (Le Marchand *et al*., 2018; Raimondi *et al*., 2011) (Figure 1 Right). To better understand if this dual backbone conformation represents under-recognized conformational heterogeneity, with potential implications for MHC binding, conformational dynamics, and aggregation propensity, we employed our electron density guided AlphaFold3 method to screen for evidence of alternate conformations within the deposited electron density maps of isolated *β*_2_M crystal structures.

**Figure 1:**
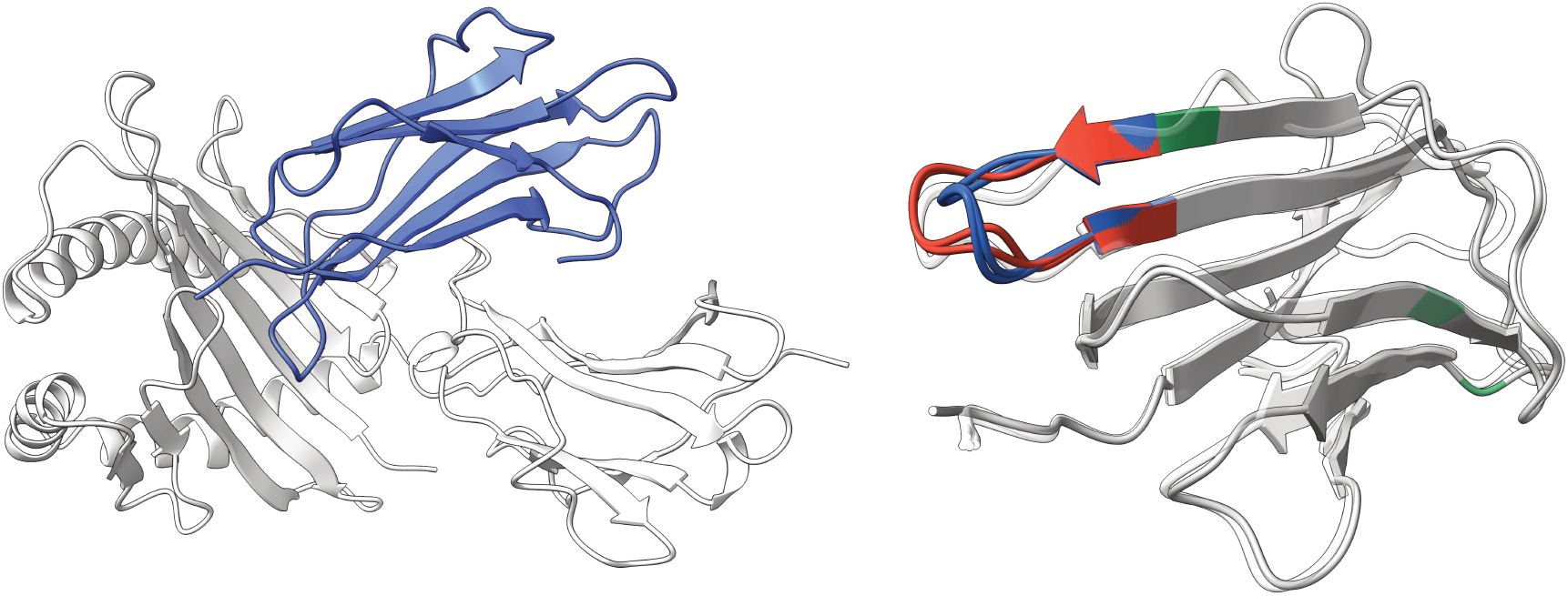
Structure of *β*_2_-Microglobulin. (Left) *β*_***2***_-Microglobulin (blue) associates with the heavy chain of the Major Histocompatibility Complex class I molecule (PDB ID 1A1M gray) to stabilize the overall fold and support formation of the peptide-binding platform. (Right) The W60 *β*_2_-Microglobulin binding loop has been modeled in alternative conformations in 4RMW and 3QDA and are shown in red (conformation A) and blue (conformation B). Mutations present across different structures are highlighted in green and AlphaFold3 prediction is overlaid in white.

## 2 Methods

### 2.1 Identification of isolated *β*_2_-Microglobulin structures for analysis

A query of the Protein Data Bank (PDB) (Burley *et al*., 2017) was performed using the RCSB PDB advanced search interface with the following criteria: (i) presence of the sequence motif `SFSKDWSFY`; (ii) polymer entity sequence length *<* 110 amino acids; (iii) a single distinct protein entity with a single polymer instance (one chain); (iv) polymer entity type restricted to protein; (v) experimental method set to X-ray diffraction; and (vi) source organism specified as *Homo sapiens*. This search returned 24 crystallographic entries, of which 22 belonging to either space group `C 121` or `I 121` were analyzed and are described in Table 1.

**Table 1:** Structures of monomeric *β*_2_-Microglobulin crystallizing in space groups `C 121` and `I 121`. For each PDB entry, the sequence mutation, resolution, polyethylene glycol (PEG) concentration used in crystallization (in %), and space group are reported. The presence of alternative conformations at the binding loop (`SFSKDWSFY`) is indicated for both the PDB structure and the electron density-guided AlphaFold3 models. Cosine similarity between observed and calculated real-space density maps carved 5Å around the sequence motif `SFSKDWSFY`, together with *R*_work_ and *R*_free_, are reported for the guided models and the deposited structures. Structures crystallized in `C 121` consistently exhibit higher map-model agreement and more frequent detection of alternative conformations compared with structures in `I 121`. Entries in **bold** indicate the superior metric between the guided and PDB entry.

| PDB ID | Variant | Res. (Å) | PEG (%) | Space group | Altloc detected | | Cosine Sim. | | $R_{\text{work}}$ | | $R_{\text{free}}$ | |
| --- | --- | --- | --- | --- | --- | --- | --- | --- | --- | --- | --- | --- |
|  |  |  |  |  | Guided | PDB | Guided | PDB | Guided | PDB | Guided | PDB |
| 2YXF | WT | 1.13 | 25 | C 121 | ✓ | ✗ | 0.881 | **0.902** | 0.180 | **0.173** | 0.185 | **0.182** |
| 4FXL | D77N | 1.40 | 18 | C 121 | ✓ | ✗ | **0.934** | 0.913 | **0.183** | 0.183 | **0.219** | 0.219 |
| 4RMW | D77A | 1.40 | 21 | C 121 | ✓ | ✓ | **0.914** | 0.906 | 0.188 | **0.187** | **0.216** | 0.217 |
| 4RMU | D77E | 1.40 | 24 | C 121 | ✓ | ✗ | **0.935** | 0.912 | **0.167** | 0.168 | **0.190** | 0.194 |
| 4RMV | D77H | 1.46 | 30 | C 121 | ✓ | ✗ | **0.929** | 0.909 | **0.193** | 0.193 | **0.231** | 0.233 |
| 3QDA | W96L | 1.57 | 19 | C 121 | ✓ | ✓ | **0.901** | 0.893 | 0.174 | **0.172** | **0.207** | 0.208 |
| 2D4F | WT | 1.70 | 20 | C 121 | ✓ | ✗ | 0.927 | **0.928** | 0.193 | **0.184** | 0.223 | **0.218** |
| 1LDS | WT | 1.80 | 21 | C 121 | ✓ | ✗ | **0.908** | 0.907 | 0.172 | **0.163** | 0.206 | **0.204** |
| 6M1B | V28M | 1.88 | — | C 121 | ✗ | ✗ | **0.940** | 0.940 | 0.212 | **0.207** | 0.264 | **0.260** |
| 7NMR | D77S | 1.15 | 32 | I 121 | ✗ | ✗ | 0.656 | **0.669** | 0.190 | **0.185** | 0.208 | **0.202** |
| 7NMC | D77E | 1.20 | 32 | I 121 | ✗ | ✗ | 0.732 | **0.739** | 0.206 | **0.194** | 0.222 | **0.208** |
| 7NMO | D77A | 1.20 | 32 | I 121 | ✗ | ✗ | **0.687** | 0.687 | 0.178 | **0.173** | 0.199 | **0.193** |
| 7NMT | D77G | 1.20 | 32 | I 121 | ✓ | ✗ | **0.628** | 0.560 | 0.191 | **0.176** | 0.216 | **0.199** |
| 4RMT | D98N | 1.24 | 27 | I 121 | ✗ | ✗ | 0.683 | **0.702** | 0.199 | **0.189** | 0.224 | **0.213** |
| 7NN5 | D77K | 1.24 | 32 | I 121 | ✓ | ✗ | 0.707 | **0.712** | 0.245 | **0.221** | 0.285 | **0.257** |
| 7NMY | D77Y | 1.25 | 32 | I 121 | ✓ | ✗ | **0.780** | 0.746 | 0.238 | **0.220** | 0.269 | **0.255** |
| 5CSG | R97Q | 1.50 | 27 | I 121 | ✗ | ✗ | 0.750 | **0.763** | 0.204 | **0.198** | 0.241 | **0.234** |
| 7NMV | D77Q | 1.51 | 32 | I 121 | ✓ | ✗ | 0.651 | **0.687** | 0.253 | **0.221** | 0.302 | **0.262** |
| 4RMR | D39N | 1.53 | 21 | I 121 | ✓ | ✗ | 0.769 | **0.771** | 0.248 | **0.240** | 0.293 | **0.287** |
| 4RMS | D54N | 1.70 | 27 | I 121 | ✗ | ✗ | 0.685 | **0.691** | 0.219 | **0.208** | 0.266 | **0.257** |
| 5CSB | D77N | 1.72 | 30 | I 121 | ✓ | ✗ | **0.787** | 0.742 | 0.206 | **0.190** | 0.243 | **0.231** |
| 5CS7 | WT | 2.10 | 30 | I 121 | ✓ | ✗ | **0.808** | 0.784 | 0.183 | **0.175** | 0.240 | **0.231** |

### 2.2 Generating ensembles using electron density guided AlphaFold3

For each PDB entry, we first generated a 2*F*_o_ − *F*_c_ Electron Number Density (END) maps (Lang *et al*., 2014). These maps were calculated from 2*mF*_o_ − *DF*_c_ coefficients in the MTZ files and are scaled to physical units of electrons per cubic angstrom (e^−^*/*Å^3^) using `F000` with bulk-solvent contributions. To ensure consistency across structures, we aligned each map to the corresponding refined atomic model obtained from PDB-Redo (Joosten *et al*., 2014). We then used `phenix.map box`, `selection radius=10` to extract a boxed region around the model with sufficient padding. To focus the analysis on the binding-loop motif `SFSKDWSFY`, we masked the density outside a 5Å neighborhood of the motif (i.e., density at voxels farther than 5Å from any motif atom was set to zero). For each density map, the lowest 25% of density voxels were removed to focus on the high-density regime. We then evaluated the probability that the electron density exceeds a threshold *t* over the range 0 − 2.5e^−^*/*Å^3^ for each structure. Specifically, we computed the empirical cumulative distribution function (CDF) of voxel densities and evaluated the complementary probability across an evenly spaced grid of thresholds. These per-structure curves enable a consistent comparison of the high-density tails of the electron-density distributions between space-group cohorts (Figure 2D). This analysis was performed as a diagnostic characterization of the maps and was not used to guide the subsequent ensemble modeling.

**Figure 2:**
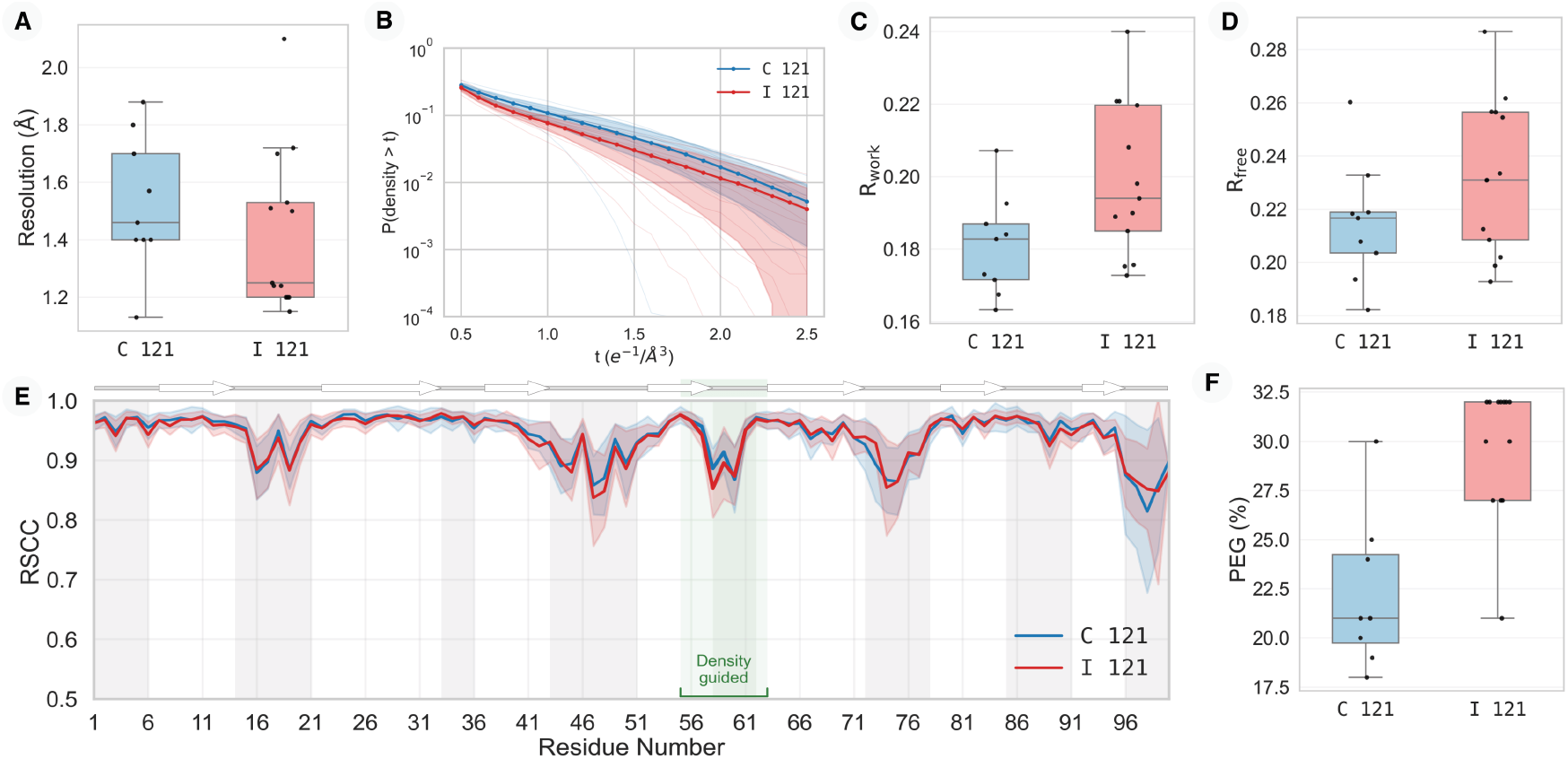
Comparison of *β*_*2*_-Microglobulin structures formed in the *C* 121 and *I* 121 space groups Despite *I* 121 crystals diffracting to superior resolution limits. **A**: and consistently yielding lower global *R*_work_ **B**: and *R*_free_ **C**: values, these bulk reciprocal-space metrics mask severe local pathologies. **D**: Log-probability distribution of electron density, *P* (density *> t*), as a function of the *σ*-threshold *t*. The *I* 121 lattice displays a rapid exponential decay at higher thresholds (*t >* 1.5), indicating a systematic disappearance of defined electron density peaks relative to the *C* 121 lattice. **E**: Residue-wise Real-Space Correlation Coefficient (RSCC). The lack of stabilizing crystal contacts in the *I* 121 lattice results in a catastrophic loss of local fit (RSCC *<* 0.6) specifically within the flexible central loop (residues 31–61), while the *C* 121 lattice stabilizes this region (RSCC *>* 0.8) through intermolecular packing. **F**: Higher PEG 4000 concentrations favour formation of *I* 121 lattices.

Once carved around the motif, we fit an ensemble of 16 structural models to the resulting local density using the procedure described in (Maddipatla *et al*., 2025a). From this initial ensemble, we selected a minimal, non-redundant subset using an Orthogonal Matching Pursuit (OMP) approach (Mallat & Zhang, 1993) as implemented in (Maddipatla *et al*., 2025a), that maximizes agreement with the observed density. The performance of the selected ensemble with respect to the PDB ensemble is summarized in Table 1. For each structure in the ensemble, we computed a normalized distance and mapped it to a diverging color scale to visualize proximity to the two reference altloc states from `3QDA` and `4RMW`. Specifically, Euclidean distances to altloc A and altloc B were computed and compared to obtain a relative distance describing which state each conformation more closely resembles. This quantity was normalized to the range [−1, +1], where values approaching −1 indicate proximity to altloc A and values approaching +1 indicate proximity to altloc B. Colors in Figures 3 and 4 encode this metric: red denotes similarity to altloc A and blue denotes similarity to altloc B, with color intensity proportional to the magnitude of the normalized distance, highlighting how strongly each conformation favors one state over the other.

**Figure 3:**
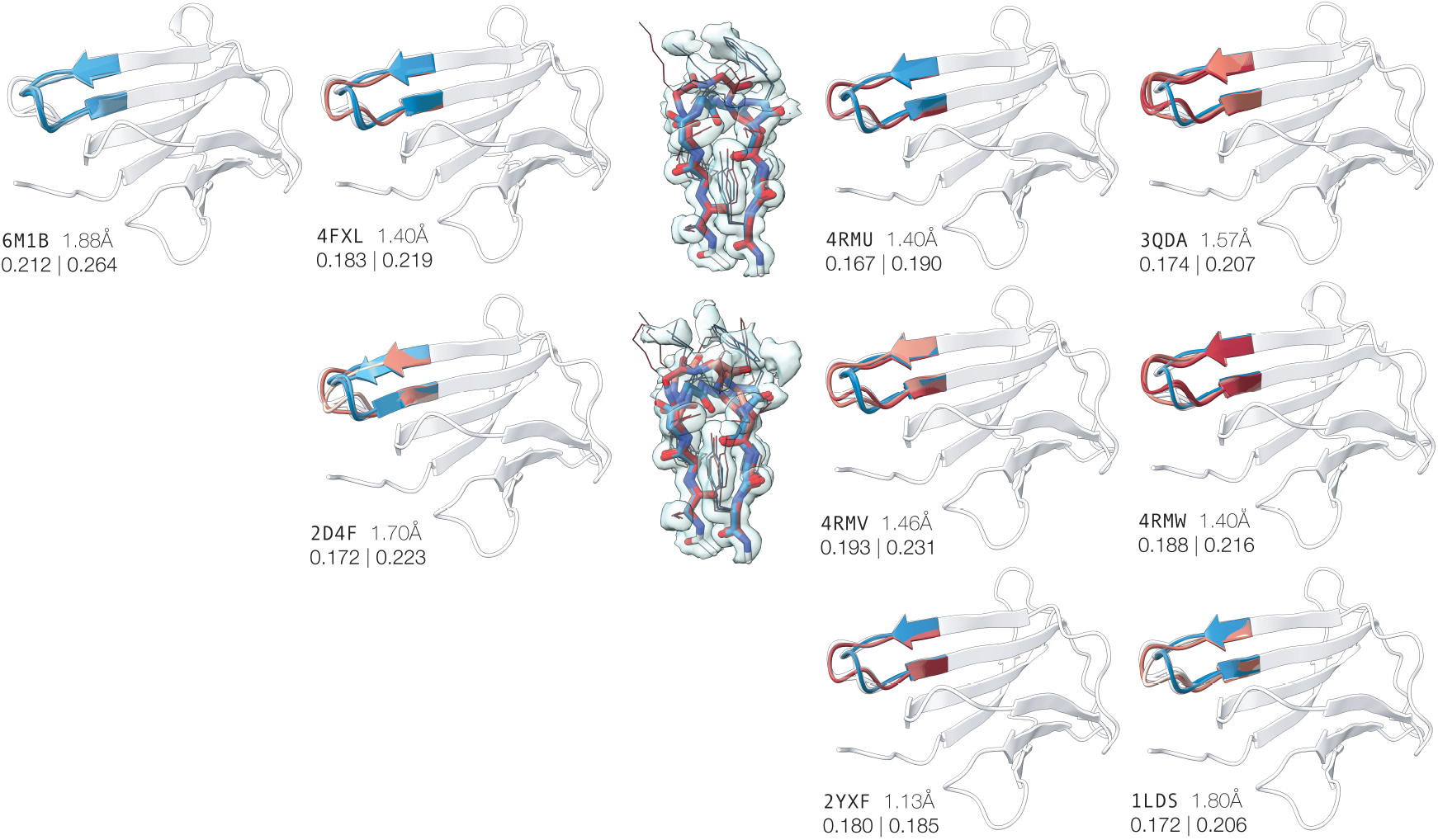
Conformational heterogeneity of the *β*_2_-Microglobulin binding loop in electron density AlphaFold3 guided models for crystal structures solved in the `C 121` space group. Each panel highlights the ensemble of conformations in the density guided loop region of each crystal, colored according to their similarity to the two experimentally observed states: conformation A (red) and conformation B (blue). The alternate conformations modeled in (`3QDA` and `4RMW`) are shown in white. The fit of ensemble to electron density (blue isosurface) is visualized for `4RMU` and `4RMV`. Panels are approximately ordered from left to right by increasing affinity to conformation A within the generated ensembles. PDB ID, resolution, *R*_work_ and *R*_free_ values for the refined ensembles are reported beneath each structure.

**Figure 4:**
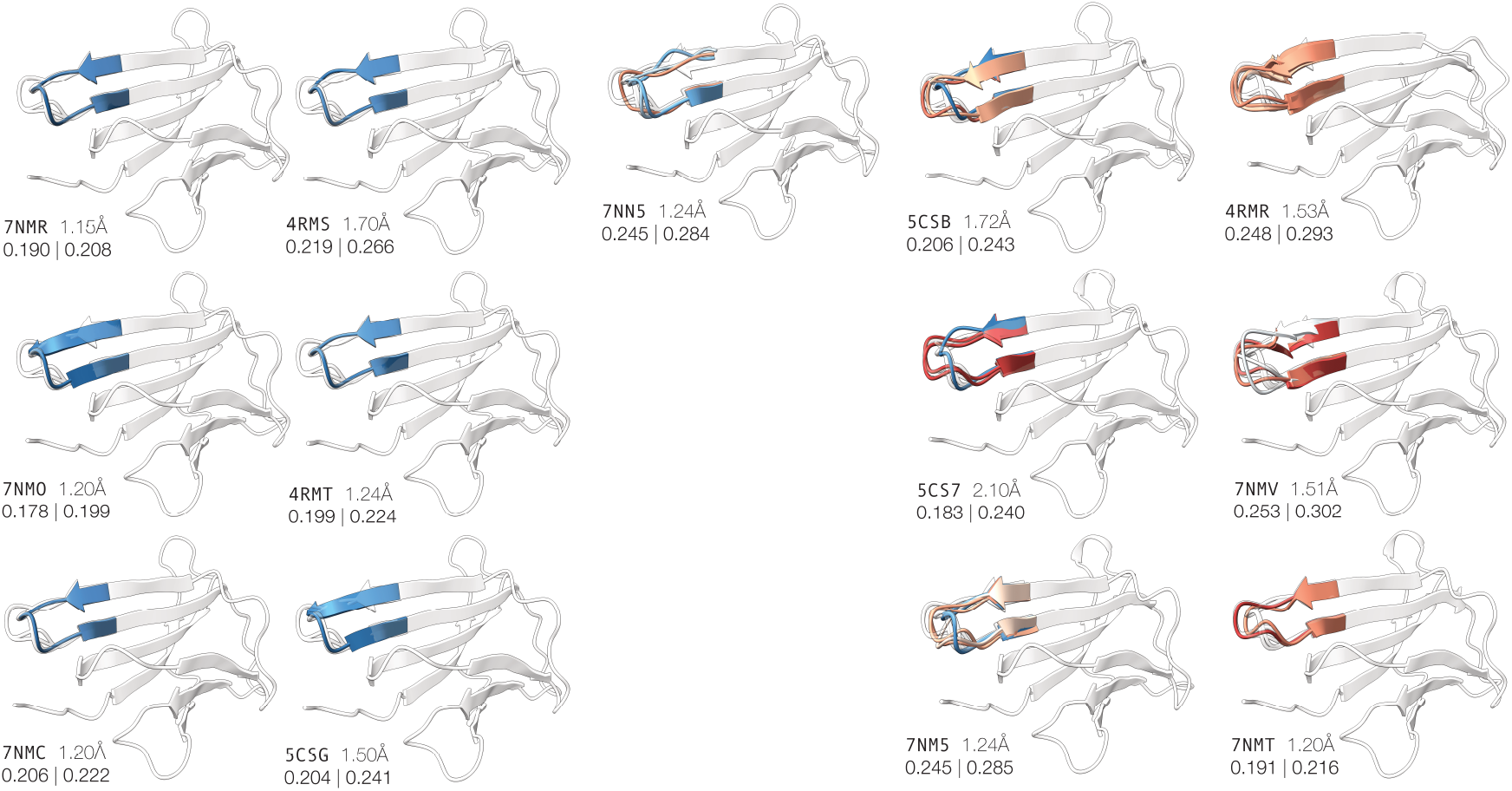
Conformational heterogeneity of the *β*_2_-microglobulin binding loop in electron density AlphaFold3 guided models for crystal structures solved in the `I 121` space group. Each panel highlights the ensemble of conformations in the density guided loop region of each crystal, colored according to their similarity to the two experimentally observed states: conformation A (red) and conformation B (blue). The alternate conformations modeled in (`3QDA` and `4RMW`) are shown in white. Panels are approximately ordered from left to right by increasing affinity to conformation A within the generated ensembles. PDB ID, resolution, *R*_work_ and *R*_free_ values for the refined ensembles are reported beneath each structure.

## 3 Results and Discussion

Compilation of monomeric *β*_2_M structures revealed that 22 of the 24 crystals segregate into two well-defined monoclinic unit-cell groups corresponding to the `C 121` and `I 121` space groups. This segregation also corresponds to the concentration of Polyethylene Glycol (PEG) 4000 used during crystallization; higher concentrations associating with the `I 121` space group. All crystals were obtained in similar crystallization conditions using PEG 4000 as the main crystallant and typically including 0.2M ammonium acetate, 15 − 25% glycerol and 0.1M acidic buffers. This grouping was also strikingly consistent with differences in the cosine similarity between the model and the density over the 5Å region carved for electron density guided AlphaFold3 ensemble generation (Table 1). A poorer fit is consistently observed for the `I 121` space group. Figure 2 demonstrates that despite having better resolutions, `I 121` group of structures have worse *R*_work_*/R*_free_ metrics and lower residue specific real space correlation coefficients (RSCC) in loop regions including the `W60` binding loop. Importantly, it is clear that these differences are not associated with variant type since in several instances the same mutation type is found in both groups (`WT`, `D77N`, `D77A`, and `D77E`).

Analysis of unit cell parameters show that the `C 121` crystals consistently form with *a* ≈ 77Å, *b* ≈ 29Å, and *c* ≈ 55Å, and an obtuse monoclinic angle *β* between 121 − 127^°^. `I 121` crystals display a similar *b* axis of ∼ 29Å but smaller and more varied *a* axes (*a* ≈ 53 − 58Å) and larger and more varied *c* axes (*c* ≈ 62 − 69Å), with a monoclinic angle (*β* ≈ 96 − 103^°^). Although the two crystal forms exhibit comparable unit-cell volumes (≈ 1.0 × 10^5^ Å^3^), they are crystallographically non-isomorphous (see Table 2). `C 121` is C-centered and lower symmetry, potentially leading to more independent reflections and better averaging, while `I 121` is body-centered and higher symmetry, potentially reducing the number of independent reflections and resulting in weaker or less interpretable density.

**Table 2:**
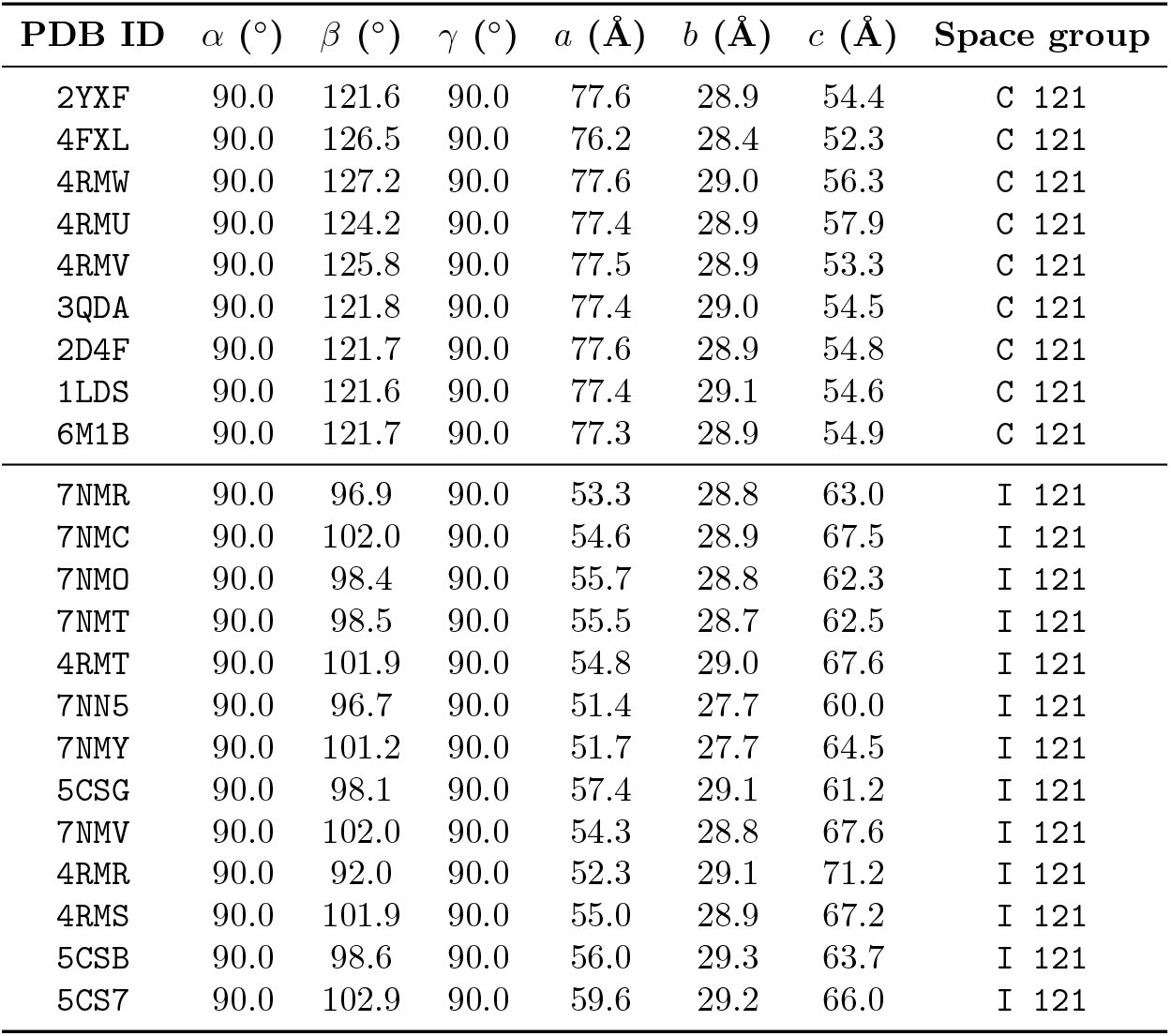
Unit-cell parameters for *β*_2_-Microglobulin crystal structures in space groups `C 121` and `I 121`, highlighting the distinct clustering of lattice dimensions and monoclinic angle *β* between the two crystal forms.

Interestingly, higher PEG 4000 concentrations preferentially induced the formation of the `I 121` lattice underscoring the importance of optimizing the phase diagram to determine crystallization conditions yielding the most informative (sets of) crystal structures (Rupp, 2015; Hashizume *et al*., 2020).

We remodeled the density for all crystals listed in Table 1 using our electron density guided AlphaFold3 method described in (Maddipatla *et al*., 2025a) and briefly outlined in the methods section. For eight out of nine crystals belonging to the `C 121` space group we detected ensembles having conformations corresponding to both of the alternately located backbones observed and modeled in `4RMW` and `3QDA` (Figure 3). The cosine similarity between the model and the density measured in the guided region, as well as the general *R*_work_ and *R*_free_, is comparable between the guided and PDB deposited models in all cases (Table 1). For two crystals we observed that the guided model gave a superior fit to the electron density as judged by improved metrics for both local cosine similarity and global *R*_work_ and *R*_free_ (Figure 3). A third crystal (4FXL) gave an equivalent fit to the PDB model (Table 1). In contrast, remodeling the density for crystals belonging to the I 121 space group often generated unimodal models and detection of both alternately located backbone conformations was observed only in about half the cases (Figure 4). As in the case of the C 121 crystals the newly generated of models had cosine similarity and *R*_work_ and *R*_free_ values comparable to the PDB deposited models (Table 1). The inconsistency between the `C 121` and `I 121` group with respect to detection of binding loop heterogeneity included comparisons of the same protein variants (`D77E` and `D77A`). This highlights a dual challenge: (i) improving ensemble modeling methods to capture minor conformations across structures of variable map quality, and (ii) recognizing how crystallization conditions influence subtle structural features. Gone are the days when a single crystal could be assumed to represent the full conformational landscape; in today’s data-rich era, multiple experimental repeats across diverse conditions are essential to systematically uncover protein flexibility and functionally relevant heterogeneity.

## 4 Conclusion

Our study demonstrates that electron density-guided AlphaFold3 provides a powerful and systematic approach to uncover hidden conformational ensembles in protein crystal structures. By re-analyzing *β*_2_-Microglobulin crystals, we revealed previously unmodelled alternative backbone conformations, particularly in the `C 121` lattice, and highlighted how lattice type and crystallization conditions influence the visibility of structural heterogeneity. These findings underscore that conventional single-conformation models often underrepresent the true conformational diversity present in crystals. Incorporating density-guided ensemble modeling allows for a more complete and accurate depiction of protein structural landscapes, offering new insights into flexibility, functionally relevant motions, and lattice-dependent effects. More broadly, this methodology provides a general framework for revisiting existing PDB structures, enhancing the interpretive power of crystallography, and bridging the gap between experimental density and the dynamic reality of protein conformations.

## Funding

Sanketh Vedula was supported in part by funding from the Eric and Wendy Schmidt Center at the Broad Institute of MIT and Harvard. Ailie Marx acknowledges the financial support of the Helmsley Fellowships Program for Sustainability and Health. Alex Bronstein is supported by the Israeli Science Foundation grant 1834/24 and ISTA HPC Cluster.

## Data availability

Generated ensembles and code for rendering the plots is available at: https://github.com/sai-advaith/b2m_actad and ensemble generation from: https://github.com/sai-advaith/guided_alphafold

## Notes

https://github.com/sai-advaith/b2m_actad

https://github.com/sai-advaith/guided_alphafold

### Competing Interest Statement

The authors have declared no competing interest.

